# Dynamics in coastal RNA viruses and bacteriophages are driven by shifts in the community phylogenetic structure

**DOI:** 10.1101/538892

**Authors:** Julia A. Gustavsen, Curtis A. Suttle

## Abstract

Marine microbes including viruses are an essential part of the marine ecosystem that forms the base of the foodweb, and drives biogeochemical cycles. Marine viral communities display repeatable changes in abundance and community composition throughout time; however, whether these changes reflect shifts in dominance within evolutionarily related groups of viruses and their hosts is unexplored. To examine these dynamics, changes in the composition and phylogenetic makeup of two ecologically important groups of viruses, and their potential hosts, were sampled every two weeks for 13 months at a coastal site in British Columbia, Canada. Changes in the taxonomic composition within DNA bacteriophages related to T4-like viruses and marnavirus-like RNA viruses infecting eukaryotic phytoplankton, as well as bacteria and eukaryotes, were examined using amplicon sequencing of gene fragments encoding the major capsid protein (*gp23*), the RNA-dependent RNA polymerase (*RdRp*) and the 16S and 18S ribosomes, respectively. The results showed that for both viral marker genes, the dominant groups of phylogenetically-related viruses shifted over time and contained many transient taxa and few persistent taxa; yet, different community structures were observed in these different viral communities. Additionally, with strong lagged correlations between viral richness and community similarity of putative hosts, the results imply that viruses influence the composition of the host communities.

**Importance:** Using high-throughput sequencing of coastal seawater collected every two weeks for one year, the dynamics of two groups of ecologically important groups of viruses were described in the context of their putative hosts and the environment. There was a large diversity of viruses and putative hosts in this study, and groups of phylogenetically-related viruses showed temporal dynamics in dominance. Examining the richness of viruses by phylogenetic groups showed different dynamics of either boom-bust or continued persistence. At the OTU-level, some members of these related groups persisted throughout time, while others were more ephemeral. These findings were put in context of potential quasispecies behaviour, and the dynamics of putative hosts. These results showed that temporal dynamics of viral communities have a phylogenetic signal which is important for understanding the ecology of these viruses since it elucidated one of the drivers of the community structure.

## Introduction

Understanding diversity, its maintenance, and its drivers is a continued theme in ecology. In microbial systems there has been extensive exploration and discussion about the mechanisms responsible for the observed high diversity (1). Many studies on microbial diversity and dynamics come from the marine milieu, where it has been argued that community composition is driven by environmental factors (2–5). Against this backdrop are viruses, which are obligate parasites that are the most abundant biological entities in the world’s oceans, and account for much of its diversity (6).

This high viral diversity arises since viruses have many different lifestyles (7), morphologies (8), and infection strategies. Some viruses infect specific strains or species of hosts, whereas others have broad host-ranges (9). As well, some groups of viruses show particularly high genetic diversity because of their low fidelity of replication (10), while others have high rates of horizontal gene transfer (11). The role of viruses as obligate pathogens, often with high host specificity, implies that they are important drivers of host composition and diversity (12); yet, our understanding of their roles as drivers of marine microbial diversity remains relatively unexplored.

Marine viruses have repeatable seasonal dynamics as revealed by measures of abundance, infectious units, and taxonomic composition. Some seasonal studies in coastal waters report that viral abundances are higher in summer than in winter (13, 14), while other multi-year time series data show that viral production and viral abundances are highest in early spring and summer (15, 16). Moreover, viral dynamics can be associated with putative hosts (16) and specific subsets of the overall coastal viral communities can show seasonal community composition dynamics (17, 18). As well, viruses infecting cyanobacteria show temporal dynamics (19, 20), with communities from the same season resembling each other more than communities sampled in the same year (21, 22), and communities being more stable in winter than in the summer and spring (23).

Viruses affect community composition in laboratory studies by reducing the abundance of the dominant host, thereby allowing others to grow up (24–26); thus, viruses promote diversity among hosts (even at the strain level) and can be responsible for large shifts in the dominant species in bacterial populations (12, 27). These dynamics have been termed “Killing the Winner” (KtW), a model in which the most actively growing hosts are killed by viruses and replaced by other strains or species (28, 29) and that co-evolution allows these dynamics to continue over time (30). There is evidence of KtW dynamics in field studies, as illustrated by a study on a solar saltern, in which coarsely-defined bacterial and viral taxa (akin to genus level grouping) were relatively constant over time, but showed KtW dynamics at a finer taxonomic scale (31). Common members of the myoviral community showed greater microdiversity over time and correlations to hosts were stronger when microdiversity within OTUs was examined in a time series in coastal California to revealing potential strain-specific effects of viruses on hosts (32). Using complete viral metagenomes recovered from a freshwater lake, Arkhipova *et al* (33) examined the dynamics of viruses over a year and found peaks in viral relative abundance before, during and after peaks in host abundance. Thus there can be interactions that do not follow the Killing the Winner model and more studies illuminating these interactions are needed as, so far, few environmental studies have examined potential Killing the Winner dynamics since few have compared hosts and viruses (34).

Examining the temporal dynamics of marine viruses and their hosts has yielded insights about the ecology of these viruses, yet little attention has been paid to the phylogenetic relationships within these communities and how they are shaped. An exception is a study by Goldsmith *et al* (35), near Bermuda, where the phylogenetic makeup of related groups of viruses over time and depth was found to be highly uneven and variable. There were differences between fall and winter attributable to stratification, with much of the variability due to one phylogenetic group of cyanophages (35). In coastal California, groups of cyanophages belonging to different phylogenetic clades shifted in their relative dominance over time (23). Knowing more about the phylogenetic diversity of the viral communities will allow us to better interpret these temporal dynamics.

Phylogenetic relatedness can be correlated to ecological relatedness in plants and animals (36, 37) and microbes have shown phylogenetic patterning in distribution and abundance (38, 39), yet little is known about these patterns in viral communities. To examine these phylogenetic patterns over time, the following hypotheses were tested: First, it was hypothesized that phylogenetic patterns in abundance would be detected in the viral communities, as has been found in putative host communities, since viruses can be driving Killing the Winner dynamics (28) or be responding to their hosts (33, 40). Second, the structure of viral communities has been purported to follow a “seed bank” distribution, where there are many more rare viral operational taxonomic units (OTUs) than abundant ones (35, 41); therefore, the ranking within phylogenetically-related viral OTUs could also follow this pattern over time. Examining the temporal relationship between phylogeny and relative abundance will reveal if genetic relatedness influences dominance in viral and, potentially, in host communities. Uncovering the alpha diversity over time in communities and how it relates to other communities and to the environment will illuminate the drivers of community structure and diversity.

To test these hypotheses, the temporal dynamics of the phylogenetic make-up of two ecologically important groups of marine viruses and their potential hosts were followed in samples taken every two weeks over thirteen months, using amplified marker genes and high-throughput sequencing. The community similarities, richness, phylogenetic diversity, and the relative abundance of phylogenetically-related groups of OTUs were examined over time in viral and putative host communities. Two groups of viruses were selected and examined because they have shown evidence of continued production in coastal waters and high diversity of viral genotypes in this environment which would be helpful in the phylogenetic patterns related to community dynamics and their diversity over time. The first group was T4-like viruses, DNA viruses that infect bacteria, including cyanobacteria. The amplification target used was *gp23*, the gene encoding the capsid (42). The second group of viruses were marnavirus-like viruses (MLVs), marine RNA viruses in the order *Picornavirales* that infect eukaryotic phytoplankton and possibly heterotrophic protists; they were targeted by amplifying part of the RNA dependent RNA polymerase (*RdRp*) gene (43). These viruses infect ecologically important phytoplankton, such as diatoms belonging to the genera *Rhizoselenia*, *Chaetoceros*, and the toxic bloom-forming raphidophyte *Heterosigma akashiwo* (44), and have been shown continued production and high diversity in oceanic communities on the west coast of North America (43, 45–47), in the North Pacific (48), in Antarctic waters (49) and in the Baltic Sea (50), and in freshwater lakes (51). Furthermore, when examined along a salinity gradient there was little overlap between the communities recovered and man unique viral OTUs recovered per site from six different sites (52). The choice of these two groups also has the advantage of encompassing groups from both DNA and RNA viruses. The dynamics of putative hosts were examined by sequencing amplified marker genes for eukaryotes (18S rRNA gene) and bacteria (16S rRNA gene). This contribution demonstrates that temporal changes in the phylogenetic make-up of viruses infecting bacteria and eukaryotic algae are related to environmental changes and to seasonal fluctuations in the communities of potential hosts.

## Materials and methods

### Sample collection

Seawater samples were collected from Jericho Pier (49° 16’36.73N, 123° 12’05.41W) in British Columbia, Canada. Jericho Pier (JP) is adjacent to the shoreline, in a well-mixed location with mixed semi-diurnal tides, and significant freshwater influence from the Fraser River. To get a representative sample of water and enough material for viral extraction, 60L of water was pumped from the 1-m depth every two weeks at the daytime high tide between June 2010 and July 2011 (33 samples). Salinity and temperature were measured using a YSI probe (Yellow Springs, Ohio, USA). For all samples, the water was pre-filtered through a 65-*μ*m Nitex mesh and filtered sequentially through 142-mm diameter, 1.2-*μ*m nominal pore-size glass-fibre (GC50 Advantec MFS, Dublin, CA., USA) and 0.22-*μ*m pore-size polyvinyldine (Millipore, Bedford, MA, USA) filters. Viral size particles in the filtrate were concentrated to ~500 mL (viral concentrate) using tangential flow ultrafiltration with a 30kDa MW prep-scale Spiral Wound TFF-6 cartridge (Millipore) (53).

### Nutrients

Phosphate, silicate and nitrate+nitrite concentrations were determined in duplicate 15-mL seawater samples filtered through 0.45 *μ*m pore-size HA filters (Millipore) and stored at −20°C until air-segmented continuous-flow analysis on a AutoAnalyzer 3 (Bran+Luebbe, Norderstedt, Germany). Chlorophyll *a* (Chl *a*) was determined in triplicate by filtering 100 mL of seawater onto 0.45 *μ*m pore-size HA filters (Millipore), and storing the filters in the dark at −20°C until acetone extraction and then analysed fluorometrically (54).

### Enumeration of bacteria and viruses

Samples for viral and bacterial abundances were taken at each sampling point by fixing duplicate cryovials containing 980*μ*L of sample with final concentration of 0.5% glutaraldehyde (EM-grade), freezing in liquid nitrogen and storing at −80°C until processing. Flow cytometry samples were processed as in Brussaard *et al* (55). Briefly, viral samples were diluted 1:10 to 1:10 000 in sterile 0.1 *μ*m filtered 1X TE, stained with SYBR Green I (Invitrogen, Waltham, MA, USA) at a final concentration of 0.5 × 10^−4^ of commercial stock, heated for 10 minutes at 80° C and then cooled in the dark for 5 minutes before processing. Bacterial samples were diluted up to 1:1000 in sterile 0.1 *μ*m filtered 1xTE, stained with SYBR Green I (Invitrogen) at a final concentration of 0.5 × 10^−4^ of commercial stock, and incubated in the dark for 15 minutes before processing. All samples were processed on a FACScalibur (Becton-Dickinson, Franklin Lakes, New Jersey, USA) with viral and bacterial samples run for 1 min at a medium or high flow rate, respectively. Event rates were kept between 100 to 1000 events per second and green fluorescence and side scatter detectors were used. Data were processed and gated using Cell-Quest software (Becton-Dickinson).

### Extraction of viral nucleic acids

The viral concentrate was filtered twice through 0.22-*μ*m pore-size Durapore PVDF filters (Millipore) in a sterile Sterivex filter unit (Millipore). Viral sized particles in the filtrate were pelleted by ultracentrifugation (Beckman-Coulter, Brea, California, USA) in a SW40 rotor at 108 000 *g* for 5 h at 12°C. The pellet was resuspended overnight in 100 *μ*L of supernatant at 4°C. To digest free DNA, the pellets were incubated with 1U *μ*L^−1^ DNAse with a final concentration 5 mM MgCl_2_ for 3 h at room temperature. Nucleic acids were extracted using a Qiamp Viral Minelute spin kit (Qiagen, Hilden, Germany) according to the manufacturer’s directions.

### PCR amplification of T4-like virus marker gene

To target the marine T4-like virus capsid protein gene (*gp23*), PCRs were set up as in Filee *et al* (42). Briefly, each reaction mixture (final volume, 50 *μ*L) consisted of 2 *μ*L template DNA, 1x (final concentration) PCR buffer (Invitrogen, Carlsbad, California, USA), 1.5 mM MgCl_2_, 0.2 mM of each deoxynucleoside triphosphate (Bioline, London, UK), 40 pmol of MZIA1bis and 40pmol of MZIA6, and 1 U Platinum Taq DNA polymerase (Invitrogen) and was amplified using the program conditions as in Table 1.

**Table 1:** PCR parameters used in this study.

| Marker gene | Target | Primer names | PCR initial | PCR denaturation | Annealing temperature | Extension | Cycles | Final extension | Reference |
| --- | --- | --- | --- | --- | --- | --- | --- | --- | --- |
| **gp23** | T4-like virus | MZIA1bis and MZIA6 | 94°C for 90 s | 94°C for 45 s | 50°C | 72°C for 45 s | 35 | 5 min at 72°C | 42 |
| **RdRp** | Marnavirus-like viruses | MPL-2F and MPL-2R | 94°C for 75 s | 94°C for 45 s | 43°C | 72°C for 60 s | 40 | 9 min at 72°C | 43 |
| **18S rRNA gene** | Eukaryotes | Euk1209f and Uni1392r | 94°C for 75 s | 94°C for 1 min | 65°C touchdown for 10 cycles followed by 55°C | 72°C for 60 s | 10 + 20 | 9 min at 72°C | 57 |
| **16S rRNA gene** | Bacteria | 341F and 907R | 94°C for 75 s | 94°C for 1 min | 64°C, 12cycles followed by 54°C | 72°C for 60 s | 12 + 25 | 10 min at 72°C | 59 ; 58 |

### PCR amplification of marnavirus-like marker gene

Half of each viral extract was used to synthesize complementary DNA (cDNA). To remove DNA, the extracted viral pellets digested with amplification-grade DNase 1 (Invitrogen). The reaction was terminated by adding 2.5 mM EDTA (final concentration) and incubating for 10 min at 65°C. Complementary DNA (cDNA) was generated using Superscript III reverse transcriptase (Invitrogen) with random hexamers (50 ng *μ*L^−1^) as per the manufacturer.

PCR was performed using primer set MPL-2 to target the *RdRp* of marnavirus-like viruses (43). Each reaction mixture (final volume, 50 *μ*L) consisted of 50 ng of cDNA, 1x (final concentration) PCR buffer, 2 mM MgCl_2_, 0.2 mM of each deoxynucleoside triphosphate, 1 *μ*M of each primer, and 1 U Platinum Taq DNA polymerase. The reaction was run in a PCR Express thermocycler (Hybaid, Ashford, UK) with program conditions as in Table 1. Products were run on a 0.5X TBE 1% low melt gel, excised and extracted using Zymoclean Gel DNA Recovery Kit (Zymo, Irvine, California, USA) as per the manufacturer and a final elution step of 2×10 *μ*L EB buffer (Qiagen).

### Filtration and extraction of marine bacteria and eukaryotes

One liter from each 60-L seawater sample was filtered through a 0.22-*μ*m pore-size Durapore PVDF 47-mm filter (Millipore) in a sterile Sterivex filter unit (Millipore). The filter was either stored at −20°C until extraction or immediately extracted as follows (56): Briefly, filters were aseptically cut and incubated with lysozyme (Sigma-Aldrich, St. Louis, MO, USA) at a final concentration of 1mg mL^−1^ for 2 h at 37°C. Sodium dodecyl sulfate was added at a final concentration of 0.1% (w/v) and each filter was put through three freeze-thaw cycles. Proteinase K (Qiagen) was then added to a final concentration of 100 *μ*g mL^−1^ and incubated for 1 h at 55°C. DNA was sequentially extracted using equal volumes of phenol:chloroform:isoamyl alcohol (25:24:1), and chloroform:isoamyl alcohol (24:1). DNA was precipitated by adding NaCl to a final concentration of 0.3M and by adding 2X the extract volume of ethanol. Samples were incubated at −20°C for at least 1 h and then centrifuged for 1 h at 20 000 *g* at 4°C. Extracts were washed with 70% ethanol and resuspended in 50 *μ*L EB buffer.

### PCR amplification of bacterial and eukaryotic ribosomal sequences

PCR targeting eukaryotes used primers Euk1209f and Uni1392r (57). These primers target positions 1423 to 1641 and include the variable region V8. Each reaction mixture (final volume, 50 *μ*L) consisted of 2 *μ*L template, 1x (final concentration) PCR buffer, 1.5 mM MgCl_2_, 0.2 mM of each deoxynucleoside triphosphate, 0.3 *μ*M of each primer, and 2.5 U Platinum Taq DNA polymerase. The reaction was run in a PCR Express thermocycler with program conditions as in Table 1.

PCR targeting bacteria used primers 341F (58) and 907R (59). These primers target the v3 to v5 regions. PCRs were run with the following conditions: each reaction mixture (final volume, 50 *μ*L) consisted of 2 *μ*L template, 1x (final concentration) PCR buffer, 1.5 mM MgCl_2_, 0.2 mM of each deoxynucleoside triphosphate, 0.4 μM of each primer, and 1 U Platinum Taq DNA polymerase. The reaction was run in a PCR Express thermocycler with program conditions as in Table 1.

### Sequencing library preparation

#### Construction

PCR products not requiring gel excision were purified after PCR using AMPure XP beads (Beckman Coulter) at a ratio of 1.2:1 beads:product. Cleaned products were resuspended in 30 *μ*L EB buffer (Qiagen). All products were quantified using the Picogreen dsDNA (Invitrogen) assay using Lambda DNA (Invitrogen) as a standard. Sample concentrations were read using iQ5 (Bio-Rad, Hercules, CA, USA) and CFX96 Touch systems (Bio-Rad). Pooled libraries were constructed using one of each of the amplicons at a concentration so that their molarity would be similar and the total product of the pool to be ~700-900 ng. Pooled amplicons were concentrated using AMPure XP beads (Beckman Coulter) at a ratio of 1.2:1 beads:product. NxSeq DNA sample prep kit 2 (Lucigen, Middleton, WI, USA) was used as per manufacturer’s directions with either NEXTFlex 48 barcodes (BioO, Austin, USA), NEXTflex 96 HT barcodes (BioO), or TruSeq adapters (IDT, Coralville, Iowa). Libraries were cleaned up using AMPure XP beads (Beckman Coulter) at a ratio of 0.9:1 beads:library.

#### Quantification and quality control of libraries

Libraries were checked for small fragments (primer dimers and/or adapter dimers) using a 2100 Bionanalyzer (Agilent, Santa Clara, CA, USA) with the High Sensitivity DNA kit (Agilent). The concentration of libraries was quantified using Picogreen dsDNA assay as above. The libraries were quantified and checked for amplifiable adapters using the Library Quantification DNA standards 1-6 (Kappa Biosystems, Wilmington, USA) with the SsoFast EvaGreen qPCR supermix (Bio-Rad) using 10 *μ*L EvaGreen master mix, 3 *μ*L of 0.5 *μ*M F primer, 3 *μ*L of 0.5 *μ*m R primer and 4 *μ*L of 1:1000, 1:5000 and 1:10000 dilutions of the libraries in triplicate on iQ5 (Bio-Rad) and CFX96 Touch qPCR machines. Cycling parameters were as follows: 95°C for 30s, 35 cycles of 95°C for 5s, 60°C for 30s, and the melt curve generation from 65°C to 95°C in 0.5°C steps (10s/step). Quantification from both Picogreen and qPCR assays were used to determine final pooling of all libraries before sequencing. Libraries were sequenced using 2×250bp PE Miseq (Illumina, San Diego, USA) sequencing at Génome Québec Innovation Centre at the McGill University (Montreal, QC, Canada), and 2×300bp PE Miseq (Illumina) sequencing at UBC Pharmaceutical Sciences Sequencing Centre (Vancouver, BC, Canada) and at UCLA’s Genoseq (Los Angeles, CA, USA).

### Sequence processing

Libraries were either split by the sequencing center using CASAVA (Illumina) or with Miseq Reporter software (Illumina). Sequence quality was initially examined using FastQC http://www.bioinformatics.bbsrc.ac.uk/projects/fastqc/). Contaminating sequencing adapters were removed using Trimmomatic version 0.32 (60) and the quality of the sequencing library further examined using fastx_quality (61). Libraries were further split into individual amplicons (i.e. 18S, 16S, *gp23* and *RdRp*) and then, if the expected overlap of the paired-end reads was 40bp or more, the paired reads were merged using PEAR (62). For 16S the expected overlap was right around the cut-off, therefore instead the sequences were aligned to the SILVA database and then assigned to either the forward or reverse primer based on their position. Then the forward and reverse reads were concatenated. However, the reverse primer sequenced less readily than the forward and thus much information was lost if only sequences with both forward and reverse were kept, therefore only the forward reads were used for further analysis. All sequences were then quality trimmed using Trimmomatic with the default quality settings. Sequences were aligned to known sequences (Silva 119 database (63) for 16S and 18S rRNA genes) using align.seqs in mothur 1.33.3 (64) and those not aligned were removed. Viral sequences were queried using BLAST against databases containing the gene markers of interest from viral isolates and from other environmental surveys and sequences with hits with an e-value below 10^−3^ were kept.

The 16S and 18S rRNA gene sequences were checked for chimeras using USEARCH version 8.0.1517 (65) with the Gold reference database. Unique, non-chimeric sequences were clustered at 97% similarity. Taxonomy for the 16S and 18S rRNA gene sequences was assigned using mothur (Wang-type algorithm) using the Silva 119 taxonomy (63). For the viral targets sequences were chimera-checked using USEARCH denovo and reference (65). Viral sequences were then translated using FragGeneScan 1.20 (66). Viral reads were clustered using USEARCH (65) at 95% similarity for MVL, and 95% similarity for T4-like viruses. Since OTU clustering sequence similarity cut-offs are not well-defined for viruses, percent similarities (at the protein level) were examined at 50% to 100% for both MVL and T4-like viruses to examine an appropriate level these values chosen for similarity. For gp23, genera can have 35% protein similarity (67). When the number of OTUs compared to the percent similarity at 95% the rise in the number of OTUs was still linear and above 95% the slope more quickly (indicating many more OTUs picked with each per percent similarity increase) thus this level it was still sensitive without splitting potential groups and similar levels and approach as in other studies (68, 69). For MLV it was as previously described (70) and chosen at 95% sequence similarity at the protein level. This approach is similar to another OTU-based study targetting ssRNA viruses in coral photosymbionts (71). Operational taxonomic unit (OTU) tables for all targets were constructed using USEARCH (65). Rarefaction curves were generated using vegan (72). Sequences were normalized for by date and by target using vegan’s rarefy (72). All of the initial sequence files have been deposited in NCBI’s BioProject database under ID PRJNA406940.

### Data analysis, multivariate statistics and phylogenetic analysis

Environmental parameters with missing data, because of instrument malfunction or unavailability, were mean imputed to fill in missing values. Day length data were retrieved using R package geosphere (73). Adonis was used as implemented in vegan (72) to test whether community matrices showed seasonal differences. Bray-Curtis distance matrices were constructed from the normalized OTU abundance tables. Mantel tests were performed by comparing the community distance matrices to each other and to distance matrices of environmental parameters as implemented in vegan (72).

NCBI CDD domain alignments for *RdRp* and *gp23* were retrieved and used as hidden Markov models via HMMER (74) to align translated OTUs with Clustal Omega (75). Environmental sequences for both *gp23* (17, 42, 76–81) and *RdRp* (43, 45) were retrieved from Genbank to give context to the OTUs.

Alignments were viewed and manually curated with aliview (82). Automated trimming of the alignments was done using Trimal (83). Initial phylogenetic trees were built with Fast Tree (84). Final maximum likelihood trees were generated using RAxML (85) with 1000 bootstraps, with VT with the PROTGAMMA model for the RdRp gene tree and JTT with the PROTGAMMA model for the *gp23* gene tree chosen using Prottest (86). Faith’s phylogenetic diversity (87) was calculated as implemented in picante (88). The package ggtree (89) was used for visualizing and annotating trees and all other plots were created using R (90). All scripts used for processing the data are available at: https://github.com/jooolia/phylo_temporal_jericho (doi: 10.5281/zenodo.2554772).

## Results

### Variability of environmental characteristics

Chlorophyll *a* (chl *a*) concentrations varied over time with a maximum observed concentration during a eukaryotic phytoplankton bloom (46.5 *μ*g L^−1^ in June 2011)(Fig. S1). The second highest chl *a* occurred during the annual spring bloom in late April 2011 (5.88 *μ*g L^−1^) which is mainly composed of diatoms belonging to *Thallassiosira* sp. (91, 92). The minimum chl *a* value of 0.05 *μ*g L^−1^ occurred in May and the chlorophyll levels remained below 1 *μ*g L^−1^ from September to March.

Nutrient concentrations were also highly dynamic ranging between 6.1 *μ*M to 67.3 *μ*M for silicate, from below 0.1 *μ*M to 2.3 *μ*M for phosphate and from below 0.1 *μ*M to 27.7 *μ*M for nitrate+nitrite (Fig. S1). Overall, nutrient concentrations were high and stable over winter, dipped in late April and then were followed by a large increase in silicate commencing in May.

### Variability of viral and bacterial abundance

During the time series, the viral abundance ranged from 5.41 × 10^6^ to 4.70 × 10^7^ particles mL^−1^ while the bacterial abundance was one order of magnitude lower ranging from 6.59 × 10^5^ to 4.43 × 10^6^ cells mL^−1^ (Fig. S1).

### Shared viral and microbial OTUs over time

Amplicon target reads representing marnavirus-like viruses (*RdRp*) and T4-like viruses (*gp23*) were translated into amino acids and the reads were normalized to the library with fewest reads. The 16S and 18S reads were normalized the same way using the nucleotide sequences. This resulted in 566 OTUs at 95% sequence similarity for the marnavirus-like viruses, with between 59 and 142 OTUs per timepoint with an average of 6% of these OTUs shared over time (Fig. S2). For the T4-like viruses there were a total of 1737 OTUs at 95% sequence similarity, with a minimum of 149 and a maximum of 484 OTUs per timepoint (Fig. S2). On average, 6% of the T4-like viruses OTUs were shared among all timepoints. There were 813 bacterial OTUs (97% sequence similarity) with an average of 10% shared over time. The lowest number of OTUs seen per timepoint was 84 and the highest number was 269. The phylum-level classifications showed a dominance of Proteobacteria and Bacteriodetes and Cyanobacteria showing large fluctuations (Fig. S3). In the eukaryotic community a total of 1115 OTUs (97% sequence similarity) were found with 6% shared on average, a minimum of 55 and a maximum of 298 (Fig. S2). The phylum-level classifications showed a dominance of the SAR group with Ophisthokonta, Haptophyta and Cryptophycaea making up the majority of the sequences recovered (Fig. S4). Rarefaction curves for individual samples did not flatten (“saturate”) indicating that not all possible OTUs were sequenced in these samples, but when considered together the curves saturated indicating that even if all the diversity was not captured in one sample, the overall community of OTUs was captured (Fig. S5).

### Dynamics of phylogenetically-related viral OTUs

Viral OTUs were placed into a phylogenetic context and their dynamics examined over time (Fig. 1,Fig. S6 and Fig. S7). Well-defined and well-supported phylogenetic groups (A-H) of marnavirus-like viruses (Fig. 1A and B) showed strong temporal dynamics and differed by season (Adonis R= 0.503, *P*-value= 0.001). Group H, which included isolated viruses infecting the diatoms *Chaetoceros* sp., and *Rhizoselenia* sp., were constant members of the communities although their relative abundance was highest in late November. Group A was always present and included many environmental sequences, as well as sequences from a virus that infects the raphidophyte, *Heterosigma akashiwo*; the group was most abundant between August and September. The winter months from October to February were dominated by OTUs in Group E, with a smaller contribution by Group H. The structure of the MVL OTU phylogenetic groups closely mirrored the structure of the top 20 OTUs found over time (Fig. S8), demonstrating that this community contained few dominant OTUs.

**Fig. 1:**
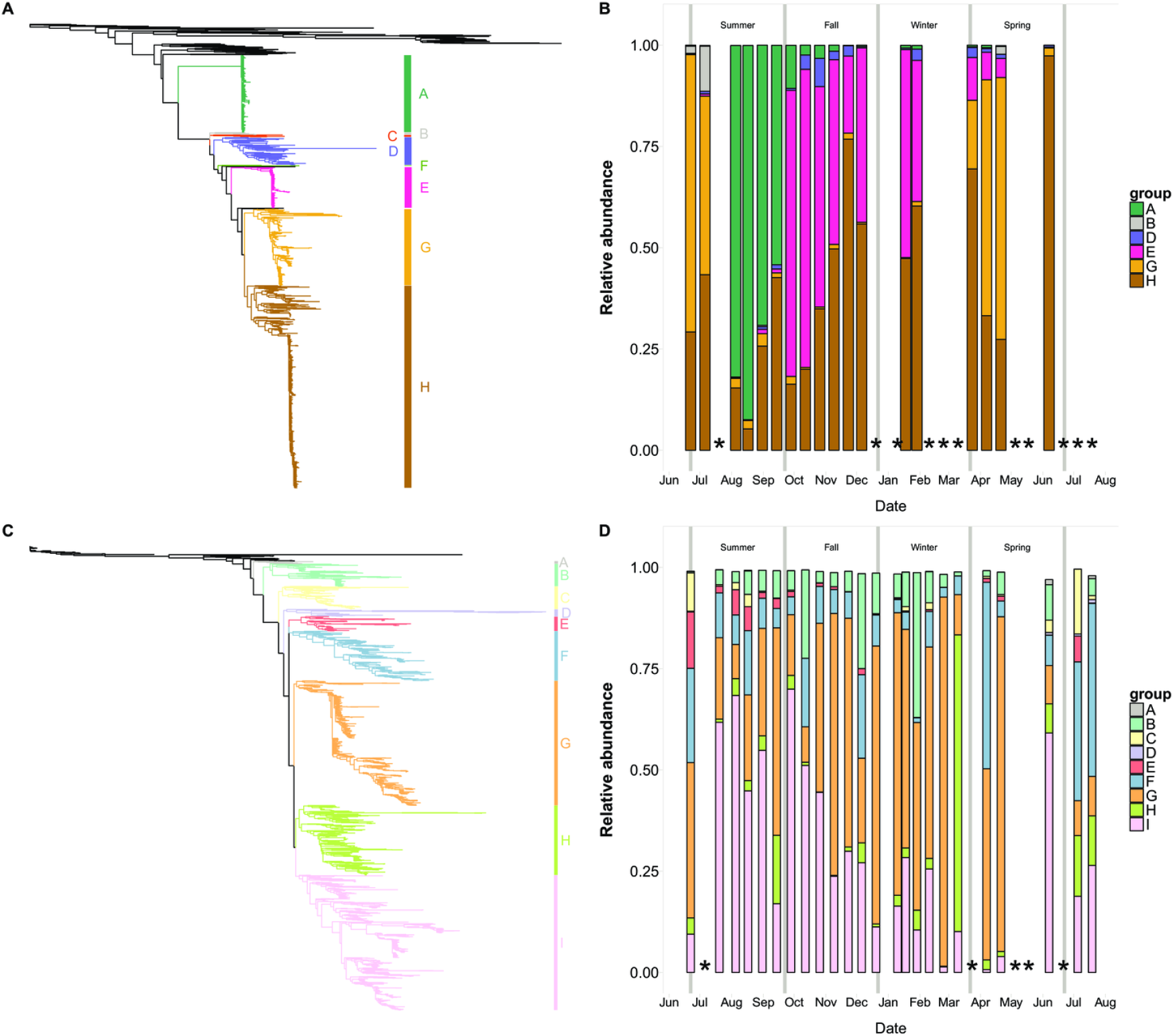
Maximum likelihood RAxML phylogenetic trees and barplots of closely-related phylogenetic groups of OTUs. A) Tree of marnavirus-like virus *RdRp* sequences including reference sequences and OTUs generated in this study. Outgroup is virus Equine rhinitis B virus (*Picornaviridae*). B) Barplot of the relative abundances of marnavirus-like virus phylogenetic groups over time. C) Tree of T4-like virus major capsid protein sequences including reference sequences and OTUs generated in this study. Outgroup is Enterobacteria phage T4. D) Barplot of the relative abundances of T4-like virus phylogenetic groups from over time. Grey vertical lines indicate seasonal boundaries. X’s indicate missing or removed samples. More detailed phylogenetic trees are available in Supplemental figures.

The T4-like virus OTUs were also placed in a phylogenetic context and categorized into groups of related OTUs (Fig. 1 C). In the fall, Group I dominated the community, followed by Group G; both groups included viral isolates infecting cyanobacteria. In January 35.7% of the relative abundance of the T4-like viruses was represented by Group B which contains no known isolates. Unlike the MVL community, the T4-like virus community had very different patterns among the top 20 OTUs and the phylogenetic groups over time (Fig. S9). When there was a large increase in nutrients in late September (Fig. S1), the dominant groups in the community shifted. The community returned to its previous state by the next sampling time. The T4-like virus communities showed small differences by season (Adonis R = 0.23, *P*-value = 0.001)

### Richness of related viruses over time

Within phylogenetic groupings the richness of OTUs varied over time in the viral communities (Fig. 2, Fig. S10 and Fig. 3, Fig. S11). Often, within the groups, the richness and relative abundance increased at the same timepoint indicating that at periods when these groups of OTUs appeared to dominate the overall communities, there was an increase in alpha diversity within the group.

**Fig. 2:**
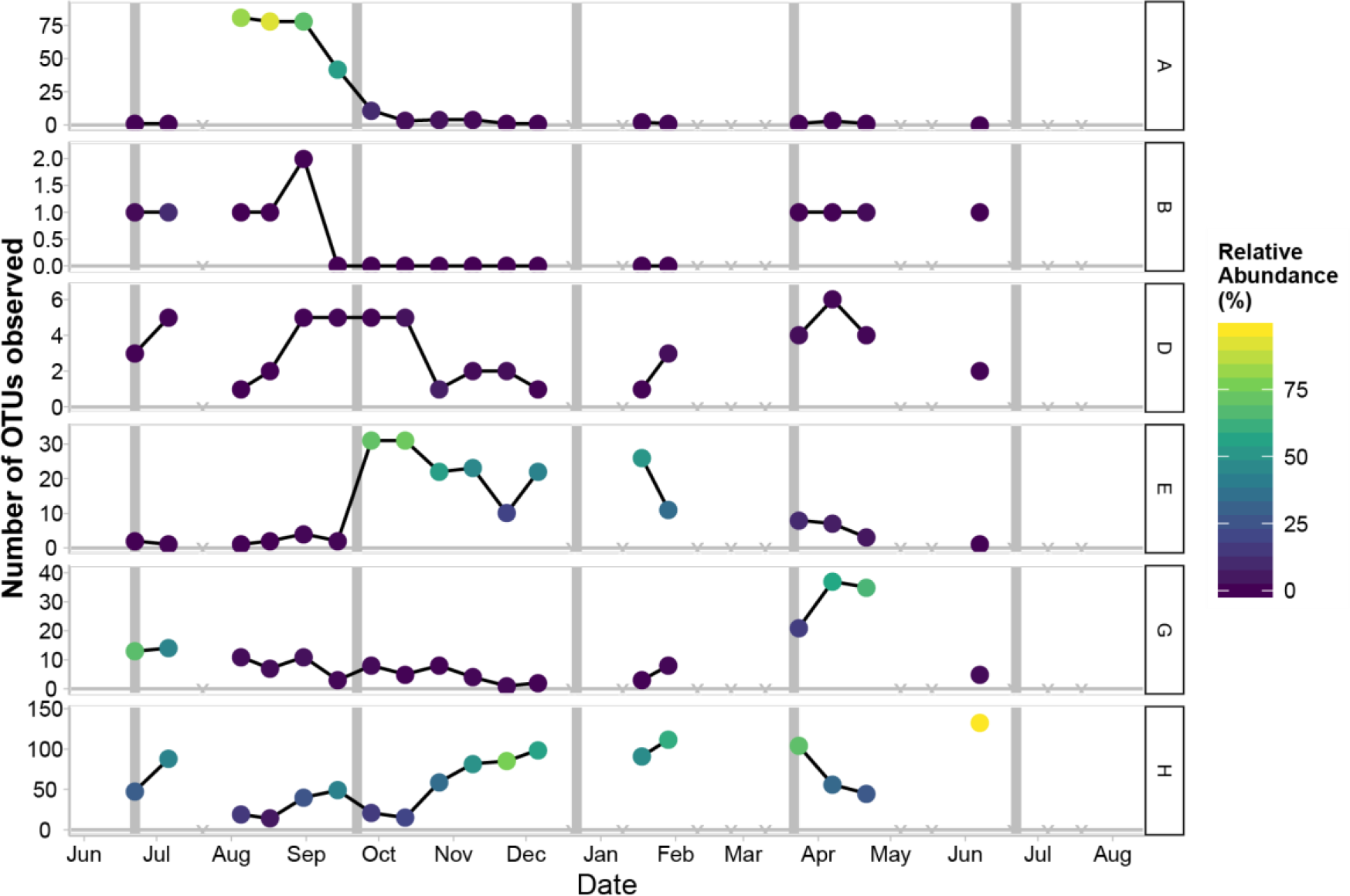
Richness of observed marnavirus-like viral OTUs (95% amino-acid similarity). X’s indicate missing or removed samples.

**Fig. 3:**
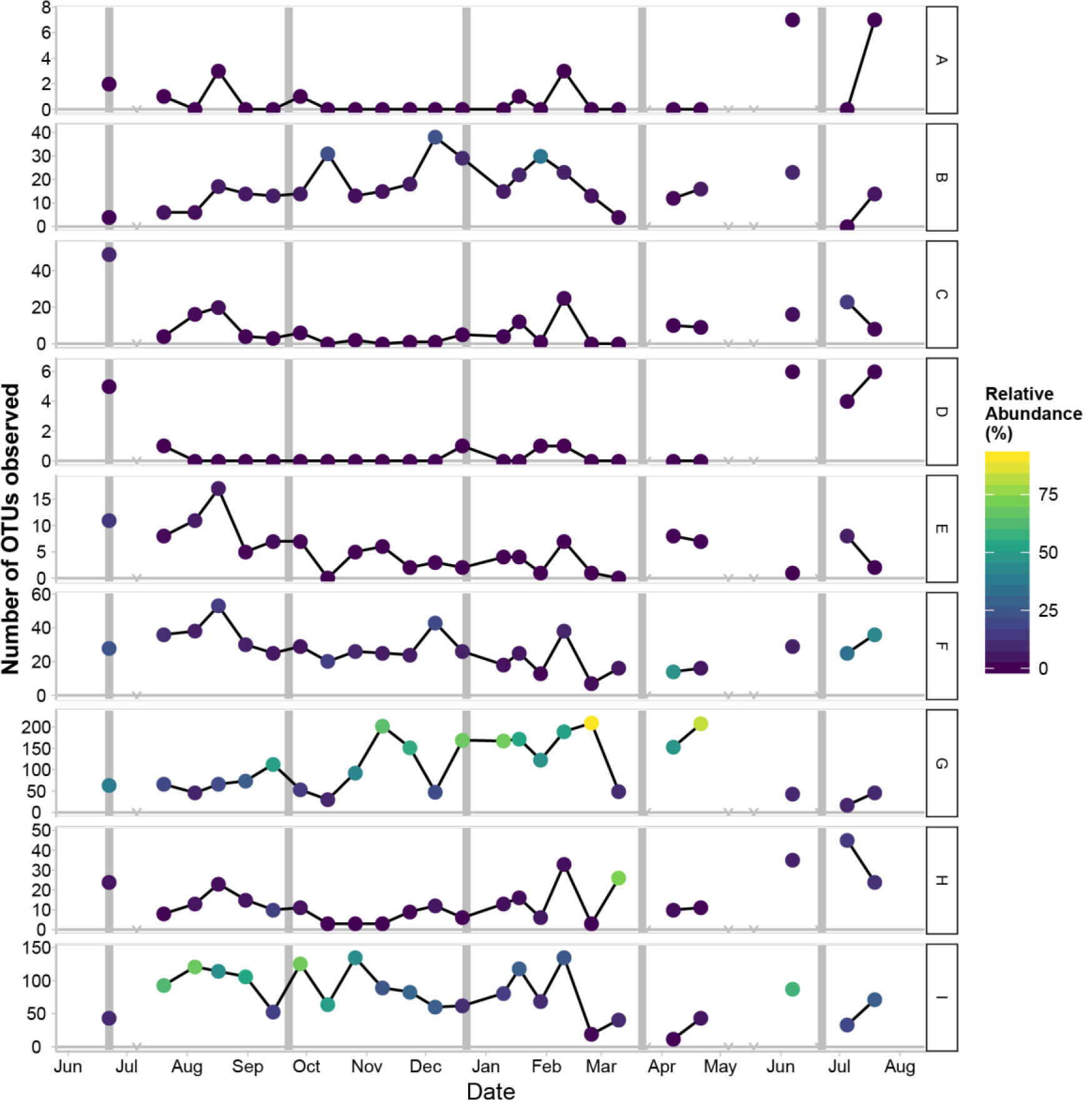
Richness of observed T4-like virus OTUs (95% amino-acid similarity). X’s indicate missing or removed samples.

Examining the richness over time for MVL viruses, for OTUs in group A there was a peak in relative abundance and richness in the Fall and later the richness dramatically decreased but this group showed continued presence over time. Similar patterns were observed for groups E and G where there was a peak in their abundance and richness at one point in the time series and otherwise a low, but detectable level of viruses. Conversely, for group H the pattern is different whereby the richness and relative abundance fluctuate but the peaks and troughs are less steep than in the other groups indicating a larger number of persistent OTUs. Group H comprised a large proportion of the overall OTUs (47% percent overall) which affects the relative dynamics observed in the MVL community.

For the T4-like viruses, Groups A, C and D had similar dynamics during the time series where they showed high richness for their group in June-August, then lower richness from August until February, and then in February there was a small peak in richness. These groups did not show large increases or decreases in the relative abundance but made up a small proportion overall of the relative abundance throughout the time series. Groups B, G, H and I had dynamics whereby they showed a continued presence throughout the time series with fluctuations between timepoints and at times a high relative abundance attributed to these groups. Groups E and F showed higher richness in the Fall than the rest of the year and a small peak in February.

There were many persistent and ephemeral OTUs over time in the viral communities (Fig. 2 and Fig. 3). Furthermore, many of the ephemeral viral OTUs were also related to the persistent viral OTUs. In the T4-like viruses there were many more OTUs than for the marnavirus-like viruses, however, both communities fluctuated over time (0.9% of the OTUs were found at 90% of the timepoints for MVL, vs. 0.8% of the OTUs in the T4-like viruses). In the T4-like viruses Groups A, B, G and F contained both persistent and ephemeral OTUs. Conversely group C had few OTUs that persisted. In the MVL viruses there were fewer OTUs overall and also fewer OTUs that persisted over time. Group A had one OTU that persisted over time and other OTUs that were only found at 4-6 sampling points, which encompassed 8 to 12 weeks at the sampling site.

### Lagged correlations with hosts over time

#### Raphidophytes and marnavirus-like viral Group A

Group A (Fig. 1A), which includes HaRNAV, a virus which infects the raphidophyte *Heterosigma akashiwo*, increased in relative abundance after the relative abundance of eukaryotic sequences classified as raphidophytes increased (Fig. 4, Spearman rank correlation over entire time series: 0.43 and *P*-value 0.07). There were further peaks in the relative abundance of raphidophyte sequences, but they did not coincide with increases in the relative abundance of marnavirus-like viral Group A. Of all the OTUs in Group A, OTU 1 was relatively most abundant, and when aligned with other closely related sequences (Fig. S12) these sequences showed changes in the amino acid D to E in the palm region of the *RdRp* (93).

**Fig. 4:**
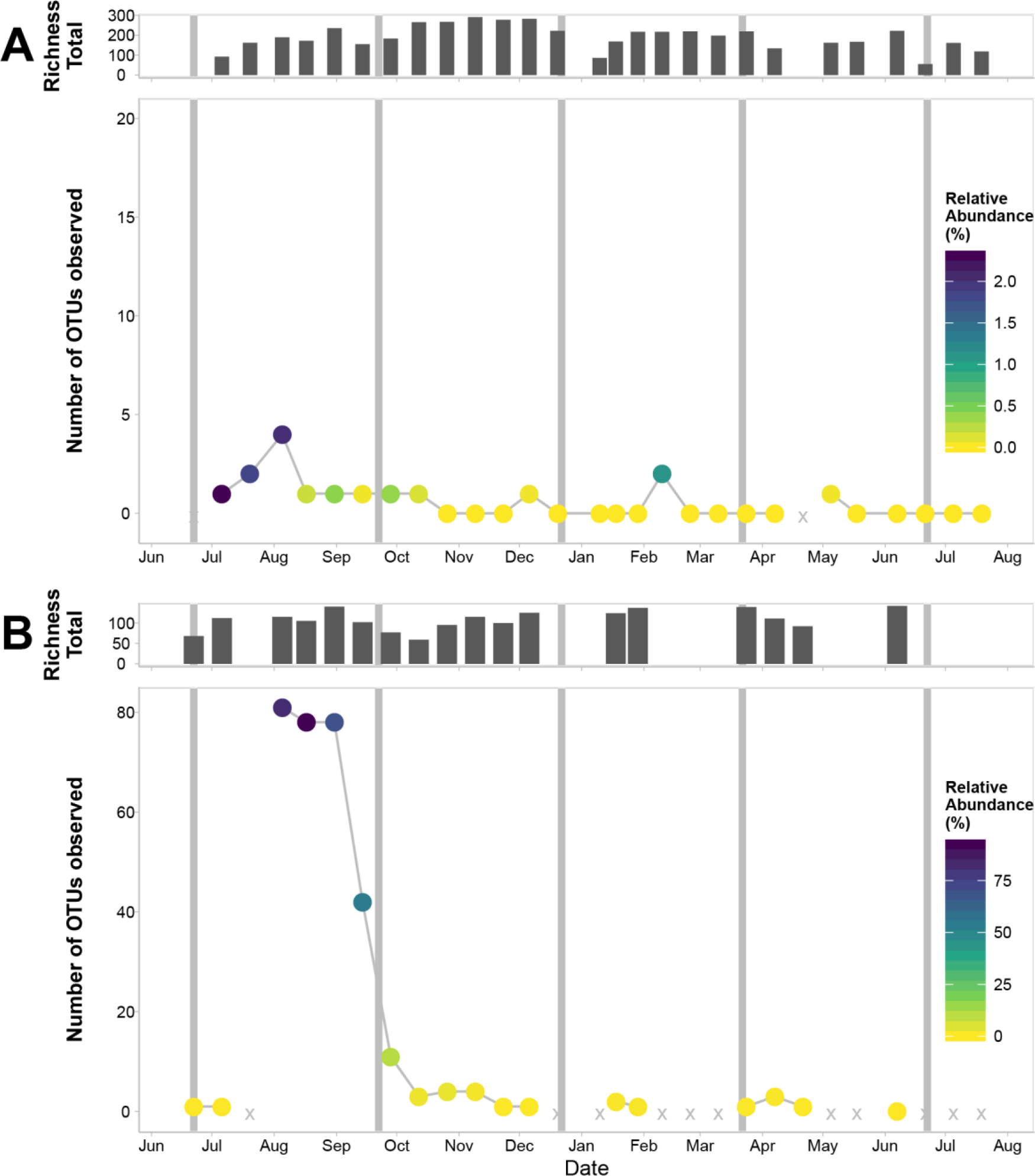
Marine marnavirus-like viral Group A compared to OTUs classified as raphidophytes over time. A) Relative abundance of OTUs (97%) classified as raphidophytic OTUs over time. Richness of all 18S OTUs at each time point is plotted above the richness of the raphidophytes.. B) Relative abundance of marnavirus-like virus Group A OTUs (95% amino acid) over time. Richness of all marnavirus-like virus OTUs is plotted above the richness of the Group A OTUs. Grey vertical lines indicate boundaries between seasons. X’s indicate missing or removed samples.

#### Cyanobacteria and T4-like virus Group I

Comparing the T4-like virus Group I, which contains cyanophage isolates, to cyanobacterial OTUs (Fig. 5), showed that the relative abundance of viruses increased in the fall after peaks in the relative abundance of cyanobacterial OTUs (correlation over entire time series: 0.56, *P*-value: 0.03). The lags in relative abundances of putative cyanophages relative to cyanobacteria continued, and after the spring diatom bloom (April 8th) there was a lag before the relative abundance of a different putative cyanophage in Group I increased, showing succession in the viral community.

**Fig. 5:**
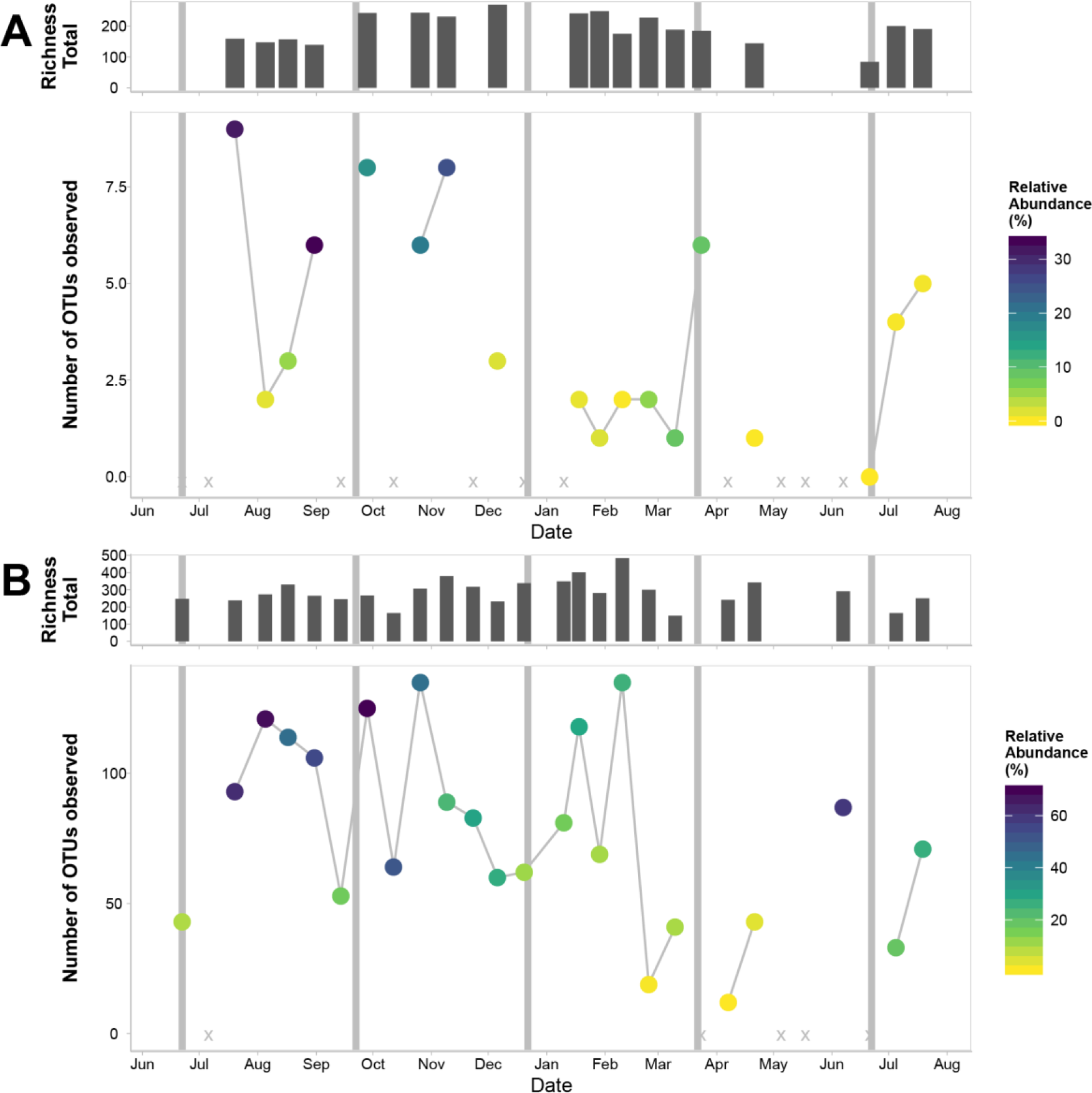
T4-like virus Group I compared to bacterial OTUs classified as cyanobacteria over time. Relative abundance of bacterial OTUs (97%) classified as cyanobacterial OTUs over time. Richness of all 16S OTUs at each time point is plotted above the richness of the cyanobacteria. Relative abundance of T4-like virus Group I OTUs (95% amino acid) over time. Richness of all T4-like virus OTUs is plotted above the richness of the Group I OTUs. Grey vertical lines indicate boundaries between seasons. X’s indicate missing or removed samples.

The community similarity of the T4-like viruses had a strong lagged negative correlation to the richness of the bacterial community (Fig. S13 and Fig. 6). The correlations to the MVL community were much stronger to other communities when lagged than when directly compared. Viral abundance was negatively correlated to bacterial community similarity.

**Fig. 6:**
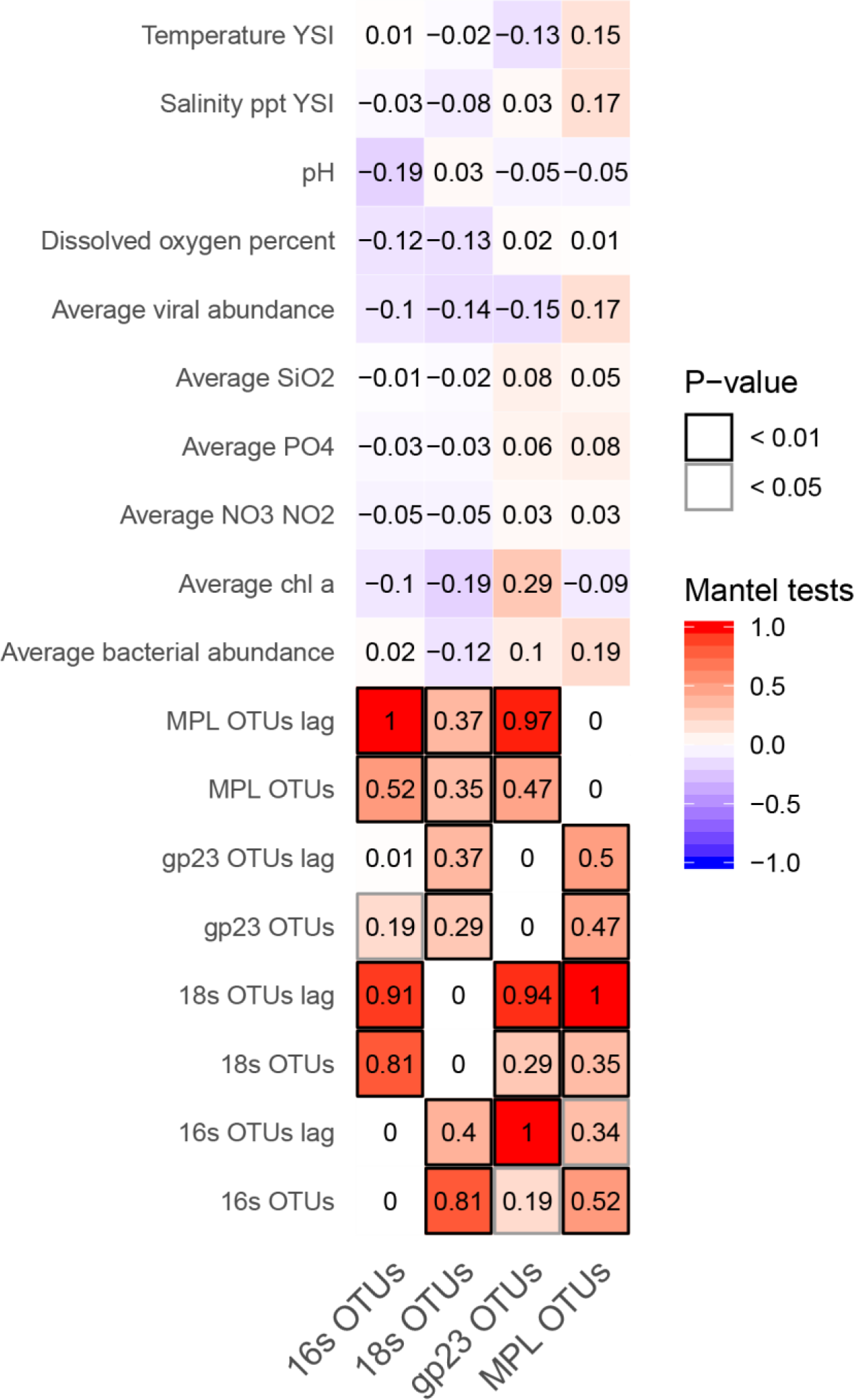
Mantel tests among community similarity matrices and distance matrices of environmental data.

#### Mantel tests among community similarity

Mantel tests were used to examine concurrent community and environmental changes over time (Fig. 6). The overall bacterial and eukaryotic community compositions fluctuated strongly together. The marnavirus-like community changes were predicted most strongly by changes in the eukaryotic community with a lag of two weeks and less strongly by the bacterial and myoviral communities. The T4-like viruses showed changes most strongly with the bacterial communities with a two-week lag, however also the eukaryotic and marnavirus-like communities with lags were also strong predictors of the myoviral communities.

## Discussion

### Temporal shifts in community dominance by groups of related viruses

A major theme in microbial ecology is understanding the causes of temporal shifts in community composition. Previously, a study at the Bermuda Atlantic Time series Study (BATS) used deep sequencing of amplicons of the viral gene marker *pho h*, to resolve phylogenetically distinct groups of viral populations that differed between fall and winter; these differences were attributed to a phylogenetic group containing cyanobacteria-infecting viruses and to water stratification (35). In our study, in coastal British Columbia, the temporal dynamics were driven by shifts in groups of phylogenetically-related OTUs (Fig. 1) and there was a consistently high diversity of OTUs in the viral communities. The dynamics of MVL viruses were similar when examined as groups of related viruses (Fig. 1B) and as the top 20 MVL OTUs over time (Fig. S8) indicating an uneven community as has been previously seen (46, 47, 70). Conversely, in the T4-like viruses the patterns among phylogenetically-related groups did not resemble the patterns of the top 20 OTUs, indicating that the community was more diverse and even. Even with these differences, the phylogenetic dynamics fluctuated in both the T4-like and MVL viruses even though viral richness stayed relatively constant over time (Fig. 2, Fig. 3,Fig. S2). Dynamics within the phylogenetic groups showed several different patterns that could be classified as having one large peak per year, several peaks or a constant presence in the community. These different dynamics could represent differences in lifestyles (r- or K-selected) (94), hosts, or the influence of environmental parameters. This would help explain how constant co-evolution or co-presence of viruses that is likely a factor in the Killing the Winner model (30).

The dynamics of the phylogenetic groups reflect that viral and microbial communities can be stable at the genus level and above, but are dynamic at the strain or species level (31). Furthermore, taxonomically related species can have similar niches and ecology (36, 37). For example, although there are exceptions, viruses infecting marine cyanobacteria and eukaryotic phytoplankton fall into different phylogenetic clusters depending on the host that they infect (95, 96), consistent with the hosts representing specific niches. Alternatively, given that the genetic similarity of marker gene sequences predicts gene content in at least some groups of double-stranded DNA viruses (96), and that viral gene content or GC ratios may result in light-dependent effects on growth or nutrient effects (97, 98), it is likely that environmental factors may have different influences on different phylogenetic groups of viruses. For example, Moniruzzaman *et al* (99) saw increased activity of ssRNA viruses with the termination of a phytoplankton bloom. Consequently, the phylogenetic signal is a very important part of the community structure.

Previously it has been found that the largest OTUs were persistent throughout time and it was suggested that transient viruses were dispersed from different areas (31). This seems unlikely in our study since it was frequent that the ephemeral viruses were closely related to the persistent viruses. Furthermore, both persistent and ephemeral viruses of phytoplankton were found in a freshwater lakes (100, 101), and at a coastal site (56), where some ephemeral viruses were correlated with shifts in environmental parameters. In another freshwater study it appeared that most viruses were transient and the system saw a very quick turnover (33). Additionally, in marine T4-like-virus communities, some OTUs persisted but many more were ephemeral in 3-year (17) and 2-year (18) time series. Furthermore, in a daily study of T4-like viral communities, few large-scale increase or decrease events were seen and overall the changes in both the bacterial and T4-like-viral communities were slow (102). Additionally, the data show that some OTUs are persistent and thus continually successful, while others are ephemeral (Fig. S10 and Fig. S11) with some of the viral groups having no persistent OTUs (T4-like C and D and MVL B, C, D, F and G). Therefore, both eukaryotic and bacterial marine viruses show the pattern of ephemeral and persistent viruses and adding the phylogenetic relatedness gives a deeper understanding of these dynamics.

### Potential quasispecies behaviour

The population structure of RNA viruses is proposed to be a mixture of genotypes called quasispecies that are produced through polymerase errors; they encompass the community of genotypes theoretically produced from one infection (103, 104). It is proposed to be a result of higher mutation rates and higher burst sizes found in RNA viruses compared to double-stranded DNA viruses (105). However, it is theoretically possible for quasispecies to exist in DNA bacteriophage populations (106). In high confidence viral DNA metagenomes there is heterogeneity in assembled reads beyond expected sequencing errors (107), and there is site-specific variation in DNA viral genome populations studied in humans (108). Thus a sequenced viral genome could be considered to be an “average” of individual genotypes. In viral RNA metagenomic sequences observed in an Antarctic lake, the ecological setting likely influences the presence of quasispecies (109) since there are more single-nucleotide variations (SNVs) in the lake-water metagenomes than in the microbial mats that they studied. This is explained either by higher turnover in the lake water, and thus more ecological niches/diversity in these samples, or as the result of convergence of water from more locations (109). When the OTUs in the marnavirus-like Group A (group which includes viral isolate *HaRNAV*) were visualized as an alignment (Fig. S12), a small number of the differences between the ephemeral and persistent OTUs were randomly spread across the gene fragment; however, most of the differences were in the “palm” section of the catalytic site C (93) where at a specific position the most persistent viral sequence had D and the ephemeral sequences had E. These ephemeral sequences point to a population that is marginally successful while the most abundant OTU (OTU_2, retaining the D amino acid) remains persistent. This suggests that the high diversity of related-viruses means a successful large-scale infection event as seen in clinical studies (104) and this suggests that this phenomenon could be prevalent in marine settings (and should be incorporated into current ecological theories (e.g. KtW)).

### Implications for theories related to community structure and dynamics

In the Killing the Winner model, viruses infect the most active organism (28). One interpretation is that hosts compete for limiting resources which determine the community composition at a particular site, and viruses determine the specific host species and the abundance of these hosts (110). With the lagged dynamics observed in the raphidophytes and viruses related to raphidophyte-infecting viruses, the “winner” has been killed (Fig. 4). Subsequently, there is a small increase in the relative abundance of raphidophytes but no associated increase in RdRp viral Group A. There are several explanations for the increase in raphidophytes without an observed increase in viral Group A. Possible explanations are that the surviving raphidophytes were resistant to specific viruses after infection (111), in this case members from viral Group A, or that the number of susceptible hosts was too low to allow for a detectable increase in viruses. Alternatively, a different subset of viruses, such as the DNA virus HAV (112), or protist grazing (113) could be keeping populations low, and thus preventing the replication of the Group A viruses. Similar patterns were detected in T4-like myoviral Group I (including isolates from Prochlorococcus phage *PSSM2* and Synechococcus phage *SSM2*) and cyanobacteria (Fig. 5). These types of patterns have been observed in other studies where there were many viral OTUs detected with strong time-lagged correlations to bacterial OTUs (40). Also, in a mesocosm experiment with *Emiliania huxleyi* there was a peak in host abundance and then four days later a peak in *Emiliania huxleyi* Virus (EhV) abundance (114). Rapid shifts in fine-scale viral dynamics and stability at coarse scale viral dynamics suggest that the Killing the Winner theory is operating at the strain level (31, 32, 115) preserving bacterial strain level diversity (12, 29).

In the marine environment, viruses, at the coarse scale of family or genus, could show Killing the Winner dynamics with finer scale dynamics at the strain level. This bank or seed bank model (41) explains how high local viral diversity (shuffling of viruses) can be consistent with low overall global diversity (represented by the most abundant viruses) by a constant local production of viruses has been supported by many studies (17, 116–118). Previously it has been found that the viral community was mostly dominated (>50%) by a few successful OTUs and the rest of the OTUs were rare and contained in the “bank” (35). We hypothesized that the communities would have a “seed bank” where there is shuffling in rank of phylogenetically-related viruses along a rank-abundance curve, however, when examined with the phylogenetic signal it is not a shuffling of rank of these related viruses as there are few viral OTUs that dominate within each phylogenetic group that dominate over time and the other OTUs are ephemeral. Thus the seed bank glosses over important information revealed by the phylogenetic examination of this communities. This seed bank idea could be effective at different time scales, but at the scale of one year with samples every two weeks there was no evidence of shuffling of rank of viruses within these viral groups. Our study has deepened this understanding by showing that the relatedness of the viruses is crucial to understanding their dynamics and reveals that the seed bank does not appear to be operating as described. Our data reveal that the viruses within phylogenetic groups can be ephemeral and related to persistent viruses and that groups of related viruses can become abundant through ecological processes (e.g. through habitat filtering where closely related species can persist in a particular environment (119)).

### Caveats

Although the challenges with viral gene markers (43, 70) and PCR in general (120) have been discussed, it remains an excellent approach for examining population structures. Although it could be argued that rare and putative quasispecies OTUs could result from PCR errors (121) or sampling anomalies (122), this is unlikely given that these OTUs were seen multiple times in different samples, implying that they are not spurious. To increase confidence in the results, some libraries with fewer reads were excluded so that more sequence could be used overall when normalizing samples (121) even though this has the trade-off of decreasing the number of samples. The sequences were checked for chimeras since chimeras can form as a result of high cycle number (123). Hence, although the read abundance of OTUs is semi-quantitative, it is a good approach for comparing richness and diversity among samples (but not for absolute counts of genes) (121). Thus, we are confident that our results conservatively reflect the changes that occurred in these microbial communities.

## Conclusions

Phylogenetically-related viruses showed temporal patterns of dominance within the viral communities over time. As well, viral communities showed evidence of time-lagged dynamics related to the potential host communities, where, for example, marnavirus-like viral (MVL) group A (containing isolate *HaRNAV*) was correlated in a lagged fashion to the raphidophytes, and myoviral virus group I (containing isolates *PSSM2* and *SSM2*) was correlated with cyanobacteria. MVL-like and myoviral viral communities differed in fine-scale community structure illustrated by differences in the proportion of OTUs that persisted over time, the evenness, and the diversity of the communities.

The MVL, which contained isolates with high burst sizes, exhibited potential quasispecies-type behaviour whereby OTUs at one timepoint were composed of many different closely related viruses and the differences between the ephemeral and persistent OTUs differed mostly in a protein residue at a catalytic site suggesting that this diversity could originate within one infection.

Previously, the link between dominance, persistence and phylogeny of virus-host communities has largely been overlooked. Other studies have found that many viral OTUs are ephemeral and that few are persistent; whereas, this study demonstrated that most of the ephemeral viral OTUs were closely related to a persistent viral OTU, and that over time the community was dominated by different phylogenetic viral groups composed of related ephemeral and persistent OTUs.

Hence, the observed changes were not the result of the presence of a seed bank that led to an overall shuffling in ranks of OTUs in the community, but rather the changes were related to fluctuations in the dominance of different phylogenetic groups of viruses over time. These dynamics thus add an important insight about the structure of viral communities and are thus crucial for understanding these viral communities.

## Acknowledgements

Suttle lab and Beaty Biodiversity members assisted greatly through discussions, manuscript improvements, assisting in field work and facilitating laboratory and analytical work. In particular the contributions of A.M. Chan, C. Chénard, C-E Chow, J.L. Clasen, C. Deeg, J.F. Finke, T.J. Heger, X.A. Tian, M. Vlok, and X. Zhong are gratefully acknowledged. C. Payne conducted the nutrient analyses and R. Pawlowicz provided the YSI probe. Thanks also to A.M. Comeau and A.I. Culley for providing reference alignments for *gp23* and *RdRp*, respectively. The research was made possible through funding from NSERC through PGS-M and PGS-D awards to JAG, and Discovery Research and Ship-time Grants to CAS, fellowships to JAG from UBC and the Beaty Biodiversity Center through a BRITE award from an NSERC CREATE grant. The Canada Foundation for Innovation and the British Columbia Knowledge Development Fund funded equipment purchases that allowed the work to be done.

